# Specialization of independently acquired flagellar FliC proteins in plant-associated *Sphingomonas* balances swimming and immunogenicity

**DOI:** 10.64898/2026.02.23.707200

**Authors:** Dor Russ, Chinmay Saha, Karnelia Paul, Zhiyu Zheng, Theresa F. Law, Manuel Anguita-Maeso, Derek S. Lundberg, Connor R. Fitzpatrick, Jeffery L. Dangl

**Affiliations:** Department of Biology, University of North Carolina at Chapel Hill, Chapel Hill, NC, USA; Howard Hughes Medical Institute, University of North Carolina at Chapel Hill, NC, USA; Swedish University of Agricultural Sciences, Uppsala, Sweden

## Abstract

Plants monitor their environment for microbial invaders using pattern-recognition receptors that detect microbe-associated molecular patterns (MAMPs). Flagellin, the main component of bacterial flagellum, contains the flg22 epitope recognized by the plant immune receptor FLS2. Immune recognition can create an evolutionary conflict, requiring bacteria to balance flagellar function and immune evasion. Here, we show that plant-associated Sphingomonads resolve this constraint by partitioning two flagellar functions, motility and colonization, across two divergent and independently acquired flagellin genes. Comparative genomics revealed widespread coexistence of FliC proteins expressing either an immunogenic variant (FliC-H) or a non-immunogenic variant (FliC-L). The non-immunogenic FliC-L is necessary and sufficient for full directional swimming, whereas FliC-H is dispensable for swimming, but sufficient for full attachment and colonization. Flagellin expression patterns mirror these functions. Thus, FLS2 recognizes the flagellar variant required for colonization rather than motility, potentially restricting colonizing bacteria from entering internal leaf and root tissues.

## Introduction

Plants coexist with diverse microbial communities that inhabit their roots and shoots (1, 2). These microorganisms range from harmful pathogens to beneficial commensals, and the lines between these lifestyles are often blurred by community contexts. Pathogens infect plant tissues to extract nutrients and proliferate, often causing damage to the host in the process. In contrast, beneficial microbes also acquire nutrients and proliferate to survive, but their growth is typically more tightly restricted by the host and constrained to specific tissues or compartments (3–5). Plants can enable microbial colonization but potentially limit beneficial microbial colonization to specific organs and compartments within them. Plants must also maintain the capacity to activate full immune responses to combat pathogens.

To monitor their environment, plants utilize plasma membrane localized extracellular pattern-recognition receptors (PRRs) that recognize conserved microbial epitopes, referred to as microbe-associated molecular patterns (MAMPs), that are typically important for microbial function (6, 7). The recognition of these microbial patterns activates MAMP-triggered immunity (MTI) (8, 9), a fundamental defense strategy that restricts pathogen invasion, but may also limit colonization by beneficial and commensal microbes, thereby fine tuning those interactions. Thus, host-associated microbial communities must reach molecular and ecological détente to ensure host health and productivity while gating the microbial community for potentially beneficial functions and against pathogens (10).

The most thoroughly characterized plant associated MAMP is flg22, a 22-amino acid peptide derived from bacterial flagellin encoded by the *FliC* gene (6, 11–14). FliC is the major structural protein that forms the bacterial flagellum (15). The FliC protein polymerizes into a helical filament that functions in bacterial motility and plays a crucial role in surface attachment, alginate acquisition, and environmental sensing (16–21). The flg22 peptide represents the minimal active epitope required to trigger immune signaling in *Arabidopsis thaliana* and many other angiosperms, but longer overlapping peptides containing the flg22 core sequence also retain immunogenicity (22, 23). Thus, immunogenicity depends primarily on the integrity and accessibility of the central flg22 region, rather than strict peptide length. Because flg22 is embedded within the intact flagellin polymer, it is not freely accessible to the plant immune system. Instead, immunogenic peptides are generated through extracellular processing of FliC in the plant apoplast (24). This involves plant glycosidases and proteases that remove glycan shields and trim neighboring peptide regions to expose or release the active MAMP, some even cleave within the flg22 epitope to relax immunity (24–26). The efficiency of active epitope release varies across microbial taxa depending on the FliC sequence and its post-translational modifications (13). The released immunogenic flg22 is then detected by the canonical plant receptor kinase FLS2 (FLAGELLIN-SENSING 2) in *A. thaliana* and other plant species, triggering a cascade of immune responses, including the production of reactive oxygen species (ROS), MAP kinase activation, and the induction of MTI transcriptional responses (6, 23). It remains unclear what part of the MTI response actually limits bacterial proliferation.

While the overall structure of flagellin is conserved, the flg22 sequence is polymorphic, and genomes from individual bacterial isolates encode diverse versions of flg22 with a range of immunogenic outcomes (11, 12, 14, 27). For example, bacteria like *Pseudomonas aeruginosa* express highly immunogenic flg22 peptides that strongly activate FLS2, whereas several commensal bacteria, including *Rhizobium*, produce variants that largely evade immune detection (11, 12, 15, 27). However, immune evasion has a cost: in the canonical flg22 variant from *P. aeruginosa,* missense mutations that evade receptor detection are also impaired for swimming (12). That trade-off is a classic case of antagonistic pleiotropy and supports a simple model where the plant immune system has evolved to target the precise amino acids and FliC structures that provide the flagellar motility function.

The flg22 antagonistic pleiotropy is further shaped by spatial selection pressures within the plant. FLS2 is not evenly distributed in the plant but rather localizes to the plasma membrane of a variety of cell types that gate access to the leaf apoplast, like stomata, hydathodes and mesophyll cells, as well as sites of lateral root emergence. FLS2 cell-type-specific expression thus positions immune perception at key microbial entry points into leaf and root tissues (28, 29). That spatial organization implies that the intensity of immune surveillance might vary across plant surfaces and organs. As a result, bacteria colonizing distinct microhabitats could experience different selective pressures to retain or modify their flagellar components. This spatial heterogeneity likely influences the evolutionary balance between flagellar functions and immune evasion, setting the stage for evolutionary innovations that might drive functional specialization and diversification of flagellar genes, as we describe below.

In this work, we surveyed a published dataset of experimentally measured immune activation of flg22 sequences from 627 plant associated bacterial genomes (11). We identified rare isolates that encode diverging *FliC* genes. Specifically, we identified six isolates of the ubiquitously plant-associated beneficial commensal genus *Sphingomonas* (30) that encode two independently acquired and divergent *FliC*-containing gene clusters. These divergent *FliC* genes encode either an immunogenic or a non-immunogenic flg22 epitope. We determined that this occurrence of duplicate *FliC* arrangement is widespread across the *Sphingomonas* sp. genomes. We focused on a single experimentally amenable isolate, *Sphingomonas* sp. MF220, and deleted each of the *FliC* genes and tested the functions of the remaining one. We found that the FliC protein carrying the non-immunogenic flg22 (FliC-L) is necessary for full swimming capacity, while the copy carrying the immunogenic flg22 (FliC-H) is sufficient for full attachment to the root and colonization of both roots and shoot. These results indicate that the plant immune receptor FLS2 targets *Sphingomonas* sp. flagella specialized in attachment and colonization, and, at least in this genus, not the classical flagellar function of motility. These results reinforce the concept that FLS2-dependent immune surveillance does not interfere with surface colonization of leaves or roots by *Sphingomonas* sp. MF220 and strains with similar *FliC* architectures but can influence their subsequent migration into intimate contact with both the leaf and the root. This restriction may contribute to partitioning of the bacteria into specific niches that allow them to thrive (and provide plant-beneficial functions) minimizing immune responses.

## Results

### Arabidopsis-Colonizing Commensal Bacterial Genomes Harbor Both Immunogenic and Non-Immunogenic Flagellin Gene Clusters

We previously presented a detailed analysis of immunogenic and non-immunogenic flg22 peptides derived from diverse FliC proteins (11, 12). We noted that some strains encoded multiple *FliC* genes. We reanalyzed this dataset that quantified the plant reactive oxygen species (ROS) burst induced by 97 synthetic flg22 epitopes from 627 *Arabidopsis*-colonizing bacterial genomes (11) to search for isolates carrying two diverging flg22 peptides: one that elicits high immune response from the plant and another that does not (Fig. 1A). We calculated the difference in immunogenicity between the most and least immunogenic flg22 epitopes for each bacterial isolate possessing multiple copies of the *FliC* gene (Fig. 1B). Most bacteria with multiple *FliC* genes, typically generated via flagellar gene cluster duplication, harbored copies with similar flg22 immunogenicity. However, eight isolates displayed flg22 copies that triggered significantly divergent ROS levels (SI Appendix, Table S1). These eight isolates contained *FliC* genes encoding both an immunogenic flg22 epitope (*FliC*-H; high ROS burst, red throughout) and a non-immunogenic flg22 epitope (*FliC*-L; low ROS burst, blue throughout) (Fig. 1B and 1C; SI Appendix, Fig S1). Six of these eight isolates belong to the genus *Sphingomonas*, a prevalent and abundant plant colonizer that expresses, among other traits, the ability to suppress pathogen infection via weak induction of host immune response (30–32).

**Figure 1.**
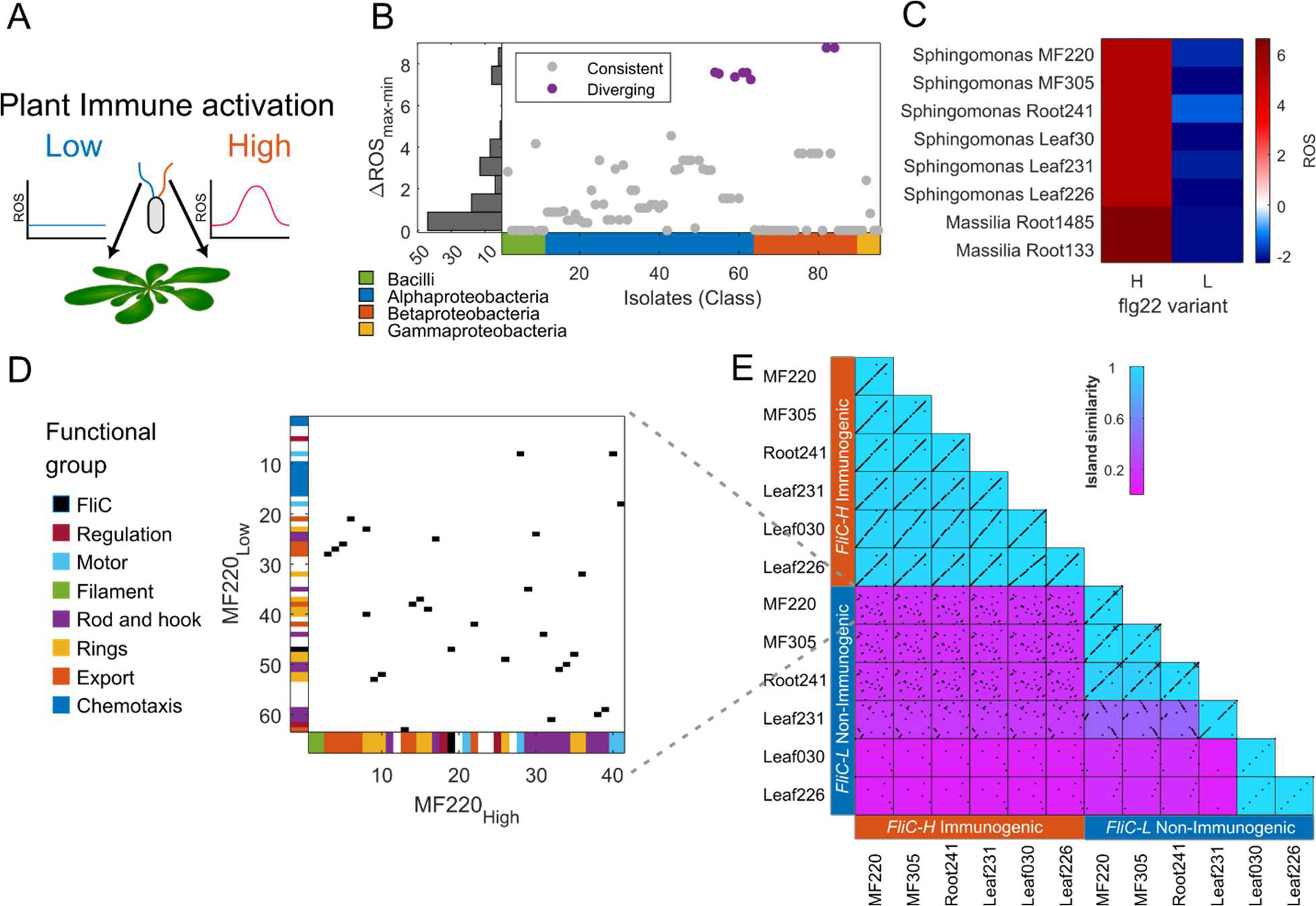
Arabidopsis-associated *Sphingomonas* encode independently evolved and acquired flagella-encoding genomic islands. **(A)** An illustration of plant-colonizing bacteria. To study the function of flagella that elicit flg22 epitope-induced immune responses (11), we defined strains that encoded two divergent flagella: (1) those expressing immunogenic flg22 epitopes (*FliC*-H, red, High, inducing ROS burst; right); (2) those expressing non-immunogenic flg22 epitopes (*FliC-*L, blue, Low, do not induce ROS burst; left). **(B)** Differential plant ROS burst between the most (max) and least (min) immunogenic flg22 variants in each Arabidopsis-associated bacterial isolate that encodes multiple *FliC* genes. Eight isolates carry *FliC* genes encoding functionally diverse flg22 epitopes (purple). **(C)** The host ROS burst elicited by the 16 flg22 peptides encoded by the eight isolates defined in **B**. ROS burst is presented as Z-score values (11). **(D)** Synteny plot of the genomic islands of the immunogenic (*FliC*-H, x-axis) and the non-immunogenic (*FliC*-L, y-axis) flagellar genes in *Sphingomonas* MF220, the reference strain for this study. Genes are colored on the axes by function. Black squares represent flagellum-related orthologous genes in the two islands. **(E)** All-by-all synteny analysis of *FliC-*anchored islands in six *Sphingomonas* isolates that encode *FliC* genes expressing both immunogenic and non-immunogenic *flg22* epitopes. Black squares represent flagellum-related orthologous genes across the two islands. Background color represents the island-similarity index, accounting for shared gene content and gene order (Methods).

We performed a synteny analysis on the genomic islands surrounding the *FliC* genes from the six *Sphingomonas* sp. strains to determine whether the two copies resulted from duplication and diversification or independent acquisition from distinct sources (Methods). We found that while the genomic islands flanking the immunogenic *FliC*-H and non-immunogenic *FliC*-L genes in *Sphingomonas* isolates contain similar flagellar biosynthetic genes, they share little similarity in gene order (Fig. 1D). Although the two islands share similar gene content (Jaccard index Ո/Ս=28/45=0.62), their distinct gene order and chromosomal locations suggest independent evolution and acquisition events. Furthermore, we observed that while the genomic islands flanking *FliC*-H are highly conserved across isolates, containing 40–42 genes, those flanking *FliC*-L are highly divergent, ranging from degenerated islands of six genes to full islands of 63 genes and defining three distinct islands (Fig. 1E; SI Appendix, Fig S2). Notably, the *FliC*-L Islands often include chemotactic machinery that is absent in all *FliC*-H islands (Fig. 1D; SI Appendix, Fig S2). The distinct gene order and accessory gene content in the genomic islands suggest that the immunogenic and non-immunogenic flagella have followed independent evolutionary trajectories and were likely acquired and maintained under distinct selective pressures.

### Plant-Associated *Sphingomonas* sp. Genomes Often Encode FliC Proteins Containing Functionally Divergent flg22 Epitopes

To characterize the prevalence of *Sphingomonas* sp. encoding FliC proteins containing functionally divergent flg22 epitopes, we analyzed a collection of 400 genome-sequenced *Sphingomonas* isolates, predominantly from geographically diverse *Arabidopsis* plants (30). We identified all *FliC* genes and the corresponding encoded flg22 epitopes (Methods). Based on sequence similarity to previously tested peptides, we defined 416 immunogenic (flg22-H, red throughout) and 143 non-immunogenic (flg22-L, blue throughout) epitopes (Fig. 2A, SI Appendix, Fig S3). Our analysis revealed that non-immunogenic flg22 epitopes exhibited greater sequence diversity than immunogenic flg22 epitopes (Fig. 2B). The sequence-diverse amino acid positions altered in the non-immunogenic flg22 epitopes include residues required for interaction with the FLS2 receptor complex (33) (Fig. 2B).

**Figure 2.**
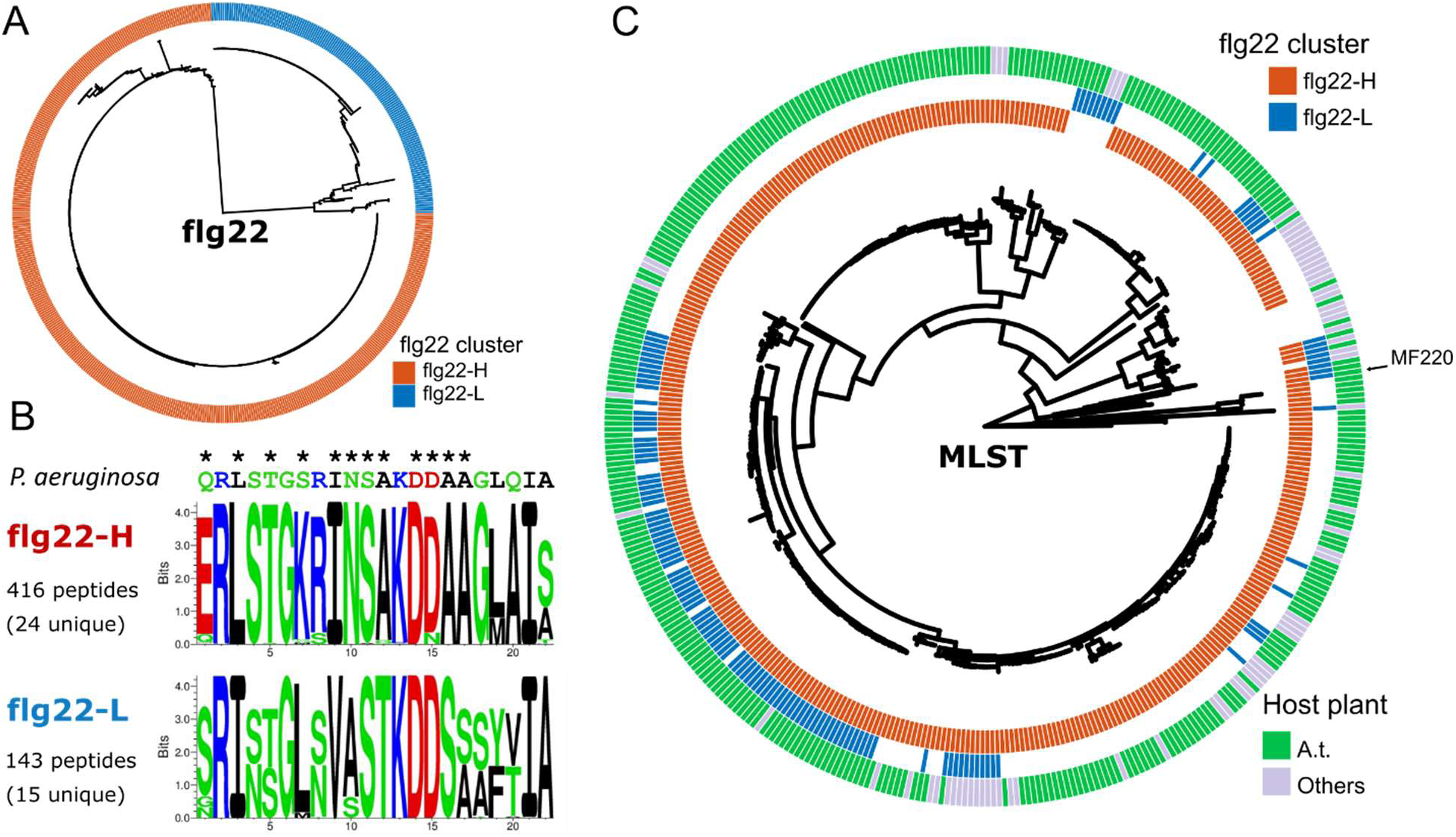
Prevalence and distribution of functionally divergent *flg22* epitopes among wild *Sphingomonas* isolates. **(A)** Maximum-likelihood phylogenetic tree of 559 flg22 epitopes derived from a collection of 400 *Sphingomonas* isolate genomes (30). The epitopes cluster into two distinct clades: those with high sequence similarity to the non-immunogenic *Sphingomonas* flg22-L epitope (blue) and those with high similarity to the immunogenic flg22-H epitope (red). **(B)** Consensus sequence motifs for the flg22-H and flg22-L epitope clades. The canonical flg22 sequence from *P. aeruginosa* appears on the top for reference (asterisks denote amino acids that contact the FLS2 receptor). Note the sequence divergence within the non-immunogenic flg22-L clade compared to flg22-H, particularly FLS2 contact residues. **(C)** Multilocus sequence typing (MLST) phylogenetic tree depicting the evolutionary relationships of all *Sphingomonas* isolates included in this analysis (MLST based on 48 conserved genes, Methods). The inner circle indicates the presence of a FliC-embedded immunogenic flg22-H epitope (red). The middle circle indicates the presence of a FliC-embedded non-immunogenic flg22-L (blue). The outer circle denotes the isolation source: *A. thaliana* (green) or other plant hosts (gray). Isolates harboring both flg22-L and flg22-H epitopes are prevalent across multiple clades and are found in bacteria isolated from both *A. thaliana* and other plant hosts.

We projected the immunogenic flg22-H and non-immunogenic flg22-L epitopes onto a multilocus sequence typing (MLST) phylogenetic tree of all isolates to characterize the evolutionary trajectories of the genomes encoding functionally divergent flg22 epitopes (Fig. 2C; Methods). The MLST tree defined the genomic diversity within this *Sphingomonas* genome collection and showed that most isolates possessed *FliC*-H genes. Isolates carrying both *FliC-*H and *FliC*-L genes were broadly distributed across different clades of the tree. A large clade (bottom-left quarter of the tree) contained many isolates with both genes; conversely, other distributed clades also contained isolates with both *FliC*-H and *FliC*-L. These results demonstrate that strains carrying *FliC* genes encoding divergent flg22 epitopes are prevalent among environmental isolates of Sphingomonas. As a whole, the divergent flg22-L epitope, the variability of the FliC-L island, and the sparse presence-absence of FliC-L across the Sphingomonas lineages suggest that FliC-L genes and islands were acquired later than FliC-H and multiple times from different sources. The widespread occurrence of isolates carrying both *FliC*-H and *FliC*-L across different clades and derived from different plant hosts suggests that this dual-flagellum strategy is a conserved feature across diverse *Sphingomonas* lineages, potentially conferring adaptive advantages in heterogeneous environments.

### The Two Diverging Flagella are Differentially Regulated and Display Distinct Morphologies

Given that the two *FliC* genomic islands likely originated from different sources, we tested whether *FliC-*H and *FliC*-L expression were co-regulated. We tested the growth of several isolates presented in Figure 2. Most did not grow well in either liquid LB or LB agar plates. Very few exhibited dispersed planktonic growth. We therefore focused on the fast-growing plant-associated isolate *Sphingomonas* MF220A (hereafter MF220). We inoculated wild-type MF220 onto 0.3% swimming agar and sampled bacteria from both the slow-swimming center of a swimming population and the fast-swimming front after three days of incubation. We measured the expression of each *FliC* gene in swimming bacteria (Methods). We found that while cells in the center of a swimming population express similar levels of FliC-H and FliC-L, bacteria at the front of the swimming halo exhibited distinctly higher expression of the non-immunogenic *FliC*-L (ratio of FliC-L/FliC-H = 189; Fig. 3A; SI Appendix, Fig. S4).

**Figure 3.**
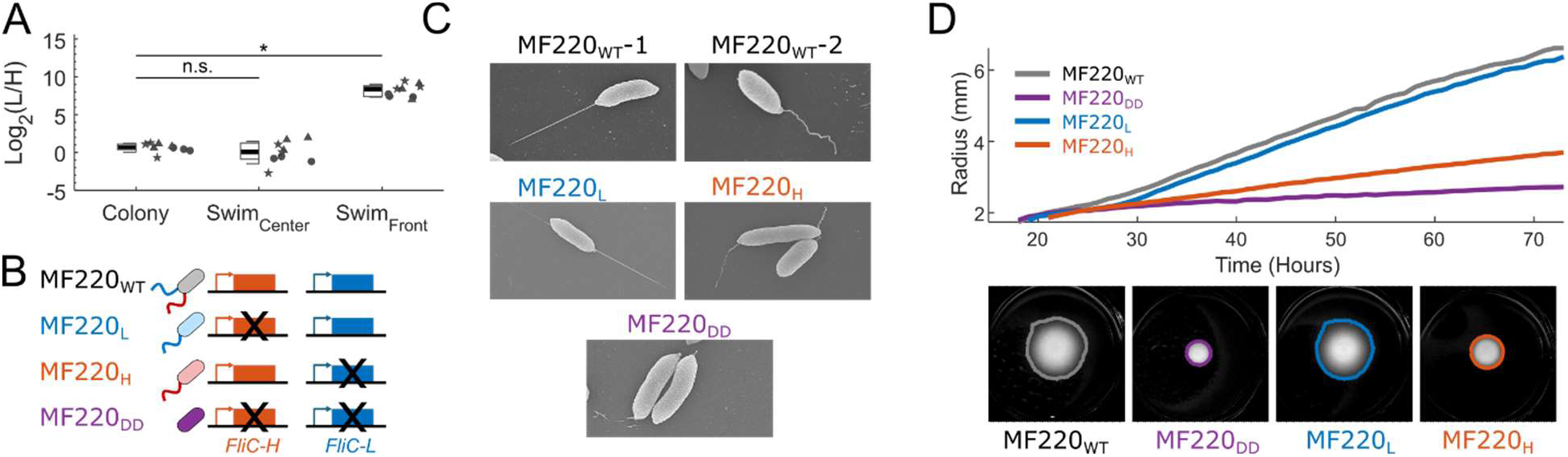
*FliC-L*, encoding a non-immunogenic flg22 epitope, is required for full swimming motility. **(A)** Expression ratio of the two *FliC* genes in MF220 from a bacterial colony on a normal 1.5% agar plate and from 0.3% swimming agar plate populations. Fast-swimming bacteria at the front of a swimming plate express 189-fold more *FliC*-L than *FliC*-H while non-swimming bacteria in a colony and from slow-swimming bacteria in the center of a swimming population express both genes to similar levels (n = 9 biologically independent samples from 3 independent experiments, denoted by shape). Boxplots show the median (black line), 1^st^ and 3^rd^ quartiles (box), and one standard deviation (whiskers) of treatment. **(B)** Illustration of the *FliC* gene deletions in *Sphingomonas* MF220. The two flagellar genes, residing in different loci on the chromosome, were deleted to generate four strains: MF220_WT_ (gray, wild-type, carrying both *FliC* genes); MF220_L_ (blue, carrying only *FliC*-L); MF220_H_ (red, carrying only *FliC*-H); and DD (purple, a double deletion lacking both *FliC* genes). **(C)** Scanning electron microscopy (SEM) of cells grown in rich media expressing either of two distinct morphologies: a straight, spear-like flagellum (observed in wild-type MF220_WT_ and in the MF220_L_ derivative) or a wobbly, twisted flagellum (observed in wild-type MF220_WT_ and in the MF220_H_ derivative). As expected, the double deletion (MF220_DD_) derivative exhibits no flagellum. **(D)** Quantification of swimming motility for MF220 derivatives on semi-solid agar. The radius of swimming populations is shown over time (Top panel). A representative final image with the identified population area is presented (bottom panel). The MF220_L_ derivative (blue), encoding only the non-immunogenic *FliC*-L gene, swims as effectively as the wild-type (MF220_WT_, gray), whereas derivatives lacking FliC-L (MF220_H_, red; and MF220_DD_, purple) exhibit severely compromised swimming.

We generated three MF220 derivatives by deleting either or both *FliC* genes to investigate their functions (Fig. 3B): the MF220_H_ derivative (red) is a deletion of *FliC*-L and thus encodes only FliC-H and thus an immunogenic flg22; the MF220_L_ derivative (blue) is a deletion of *FliC*-H and encodes only FliC-L and thus a non-immunogenic flg22; the double deletion derivative (MF220_DD_, purple), lacks both *FliC* genes (Methods). We identified two distinct wildtype MF220_WT_ polar flagellar morphologies under a scanning electron microscope (SEM) that did not obviously co-occur in the same cell: the majority of the cells carry a straight, spear like flagellum (Fig 3C, MF220_WT_-1; SI Appendix Fig. S5) while others carry a wobbly and twisted flagellum (Fig. 3C, MF220_WT_-2; SI Appendix Fig. S5). As expected, the double *FliC* deletion derivative MF220_DD_ expresses no flagella, but rather only the initial hook structure (Fig. 3C; SI Appendix Fig. S5).

Looking at the single mutant, we found that while most MF220_L_ bacteria carry the straight, spear-like flagellum, most MF220_H_ bacteria carried the twisted and wobbly flagellum (Fig. 3C; SI Appendix Fig. S5). These distinct flagellar morphologies may underline functional differences in propulsion efficiency navigation and/or host interaction, further supporting a potential division of labor between FliC-H and FliC-L in MF220.

### FliC-L is Required for Directional Swimming

We next tested the role of the two flagellar types in directional swimming along a nutrient gradient using MF220_WT_ and the three *FliC* deletion derivatives. We quantified the radius of each directionally swimming population over 72 hours (Fig. 3D; Methods; Video S1; SI Appendix, Fig. S6). We found that while the MF220_L_ derivative (blue) swam as effectively as the wild-type MF220_WT_ (black), the MF220_H_ derivative (red) exhibited severely compromised swimming. The double deletion MF220_DD_ (purple) swam more poorly than the MF220_H_ derivative (Fig. 3D). The sufficiency of the FliC-L flagellum for full directional swimming is consistent with our expression data (Fig. 3A). These results suggest that the immunogenic FliC-H flagellum is not required for directional swimming and cannot substitute FliC-L, likely because it fails to effectively engage the chemotactic machinery optimized for FliC-L mediated motility (SI Appendix, Fig. S2).

### FliC-H, Encoding an Immunogenic flg22 Epitope, is Required for Full Colonization and Attachment to *A. thaliana*

We next investigated the contribution of each of the two flagella types to plant colonization. We began by testing the first step in plant colonization, attachment. We performed short-term attachment assays (washing to eliminate loosely adherent bacteria and then grinding tissue to free tightly adherent and endophytic compartment bacteria for cfu counting; Methods) using the four MF220 derivatives, quantifying bacterial load after four hours of attachment to *A. thaliana* shoots and roots in water (Fig. 4A; Methods). In the shoot (Fig. 4A, top), the double-deletion derivative (MF220_DD_, purple) exhibited compromised attachment. By contrast, either of the two FliC-containing flagellar types was sufficient to provide wild type levels of attachment. That result suggests that while a flagellum is required for optimal attachment, the two flagellar types are redundant for shoot attachment. On roots (Fig. 4A, bottom), on the other hand, both the double-deletion derivative (MF220_DD_, purple) and MF220_L_ (blue) also exhibited compromised levels of attachment. Although our data do not resolve the individual steps of surface attachment, these results demonstrate that FliC-H (red) is sufficient for full attachment to the root, while FliC-L is not.

**Figure 4.**
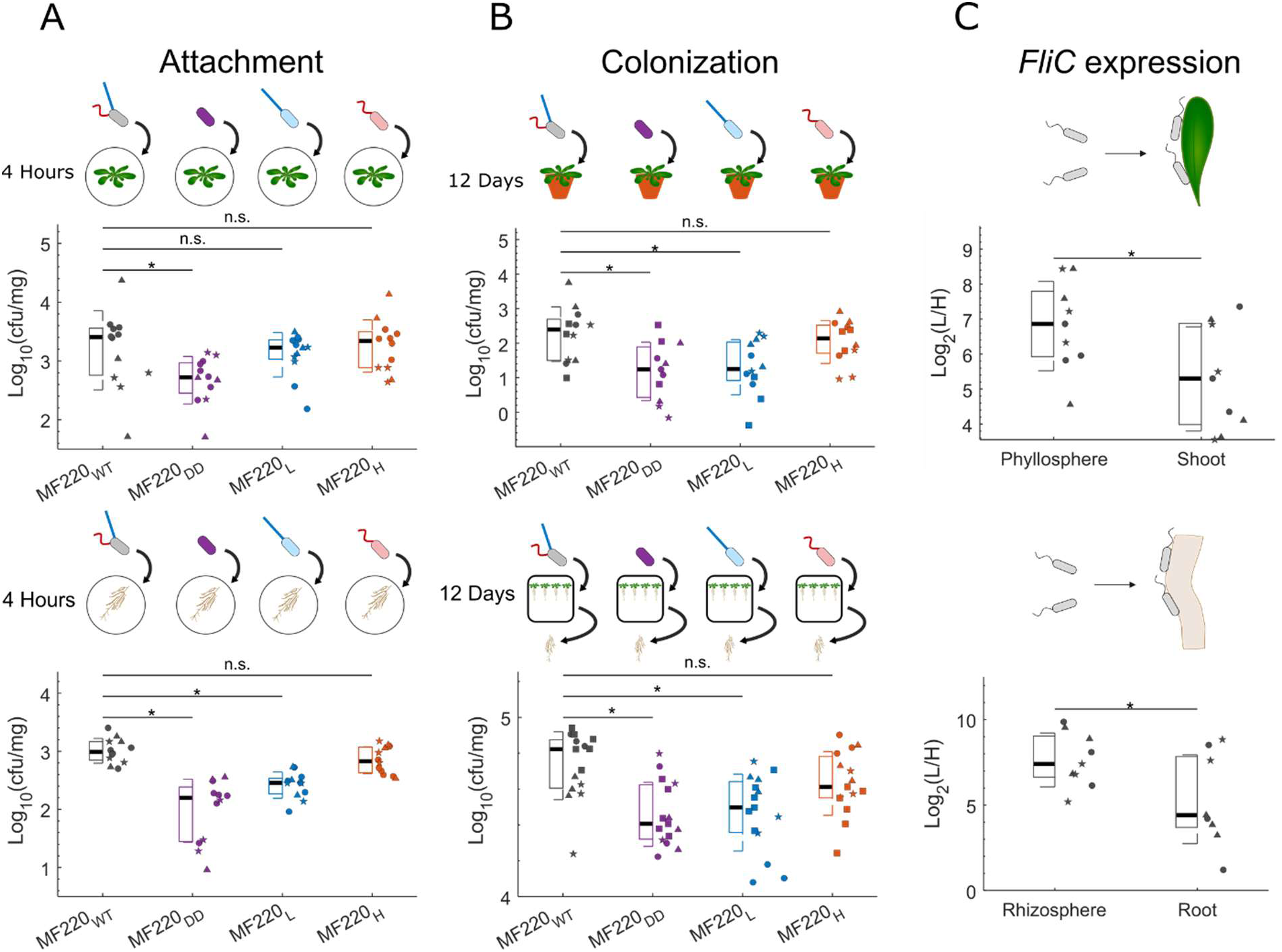
The immunogenic FliC-H is essential for robust colonization and attachment to *Arabidopsis thaliana* roots. **(A)** Quantification of short-term attachment of MF220 derivatives at four hours post-colonization to *A. thaliana* shoots (top) and roots (bottom) in liquid. MF220_DD_ (purple) bacteria were compromised in short-term attachment to *A. thaliana* shoots compared to MF220_WT_ (gray). By contrast, both the MF220_H_ (red) and MF220_L_ (blue) derivatives exhibited attachment levels similar to MF220_WT_. In the root, MF220_DD_ (purple) bacteria were again compromised in short-term attachment. Additionally, MF220_L_ also exhibited compromised attachment (linear model, *P* < 0.05, n = 12 biologically independent samples from 3 independent experiments, denoted by shape). **(B)** Quantification of MF220 derivatives colonizing the shoots (top, SI Appendix, Fig S6) and roots (bottom) of *A. thaliana* for 12 days. In both plant organs, the double deletion MF220_DD_ (purple) and the MF220_L_ (blue) derivatives exhibited significantly reduced colonization compared to the wild-type MF220_WT_ bacteria (gray). Conversely, the MF220_H_ derivative (red) colonized at MF220_WT_ levels (linear model, *P* < 0.05, n = 15 biologically independent samples from 4 independent experiments, denoted by shape). **(C)** Expression ratio of the *FliC* genes in *Sphingomonas* MF220 bacteria colonizing the shoots (top) and roots (bottom) of *A. thaliana* for 12 days analyzed by RT-qPCR. Plant-associated bacteria (right, comprising strongly attached bacteria and endophytes) exhibit significant shift in the *FliC*-L to *FliC*-H expression ratio towards *FliC*-H compared to the non-attached bacteria in the phyllosphere (top, 4.9 fold) and rhizosphere (bottom, 2.8 fold)(left; linear model, *P* < 0.05, n = 9 biologically independent samples from 3 independent experiments, denoted by shape). In all panels, boxplots show the median (black line), 1^st^ and 3^rd^ quartiles (box), and one standard deviation (whiskers) of treatment.

To assess the roles of *FliC* genes during long-term plant colonization, we tested the colonization of the four MF220 derivatives for 12 days in two systems. To test shoot colonization, we spray-inoculated *A. thaliana* leaves on plants grown in potting soil with each of the four derivatives separately (Methods). We quantified shoot colonization over 12 days by grinding washed *A. thaliana* leaves (Methods) and plating the colonizing bacteria on MF220-selective Rifampicin plates. Thus, this assay captures tightly adherent surface bacteria and those that have colonized the leaf apoplast. We found that the abundance of all four derivatives decreases over time (SI Appendix, Fig S7). Nevertheless, MF220_WT_ (black) and MF220_H_ (red) persist, and hence are more abundant than MF220_DD_ (purple) and MF220_L_ (blue) after 12 days (Fig. 4B, top). To test root colonization, we used a gnotobiotic system. We inoculated 0.5x MS plates independently with each MF220 derivative before transferring seven-day-old *A. thaliana* seedlings onto the pre-inoculated plates. After 12 additional days, we quantified the root-associated bacteria (operationally defined as tightly attached bacteria at the root rhizoplane and the root endophytes; Methods). We found that both the double deletion MF220_DD_ (purple) and the MF220_L_ derivative (blue) exhibited compromised root colonization (Fig. 4B, bottom). In contrast, the MF220_H_ derivative (red) colonized roots at levels comparable to wild-type MF220_WT_ (black). These results highlight the specific role of FliC-H in the colonization of both the roots and the shoot of *A. thaliana.* We additionally determined that these phenotypes are independent of the flg22 receptor FLS2 (SI Appendix, Fig S8).

The requirement of FliC-H for colonization led us to test flagellar expression during colonization (Methods). We compared *Sphingomonas* MF220_WT_ bacteria loosely attached to the plant tissue (root rhizosphere and leaf phyllosphere; see Methods for the methodological definitions of these compartments) (34) with those colonizing the ‘root’ (defined operationally as the combined firmly attached rhizoplane and endophytic bacteria) or the’shoot’ (defined operationally as ethanol resistant shoot endophytes). We observed that the ratio of *FliC*-L to *FliC*-H expression shifts towards greater *FliC*-H expression in both organs in those fractions associated with more intimate contact with the host (Fig. 4C; SI Appendix, Fig S4B). The shift towards higher *FliC*-H expression during plant advanced colonization is in alignment with the requirement for FliC-H in colonization (Fig. 4B).

## Discussion

Our study reveals a previously unrecognized strategy by which the prevalent and plant-enriched *Sphingomonas* sp. resolve the evolutionary conflict between flagellar function and immune evasion. We identified isolates of this genus that encode two independently acquired, divergent flagellin genes: an immunogenic (*FliC*-H) and a non-immunogenic (*FliC*-L). Further characterizing the function of each flagellar variant in an experimentally tractable isolate revealed a striking division of labor: the immunogenic variant (FliC-H) is specialized for host attachment and colonization, whereas the non-immunogenic variant (FliC-L) is specialized for directional swimming. Both contribute to short term bacterial attachment to shoots. Our data does not rule out the possibility that FliC-H promotes colonization by improving the bacterial attachment to the plant or that FliC-H is required for host surface sensing, and hence MF220_L_ bacteria are locked in planktonic phase. These two flagella differ not only in immunogenicity and function but also in their expression patterns, genomic context, and morphology, suggesting functional specialization into motility and colonization roles. Thus, rather than diversification of a single flagellin to balance motility, attachment, and immune evasion, many *Sphingomonas* may partition these functions across two distinct loci. Most importantly, our findings suggest that the plant immune receptor FLS2 recognizes the flg22 epitope derived from the *Sphingomonas* sp. flagellar variant required for intimate plant colonization rather than the flagellar variant required for motility toward and on plant surfaces (Fig. 5).

**Figure 5.**
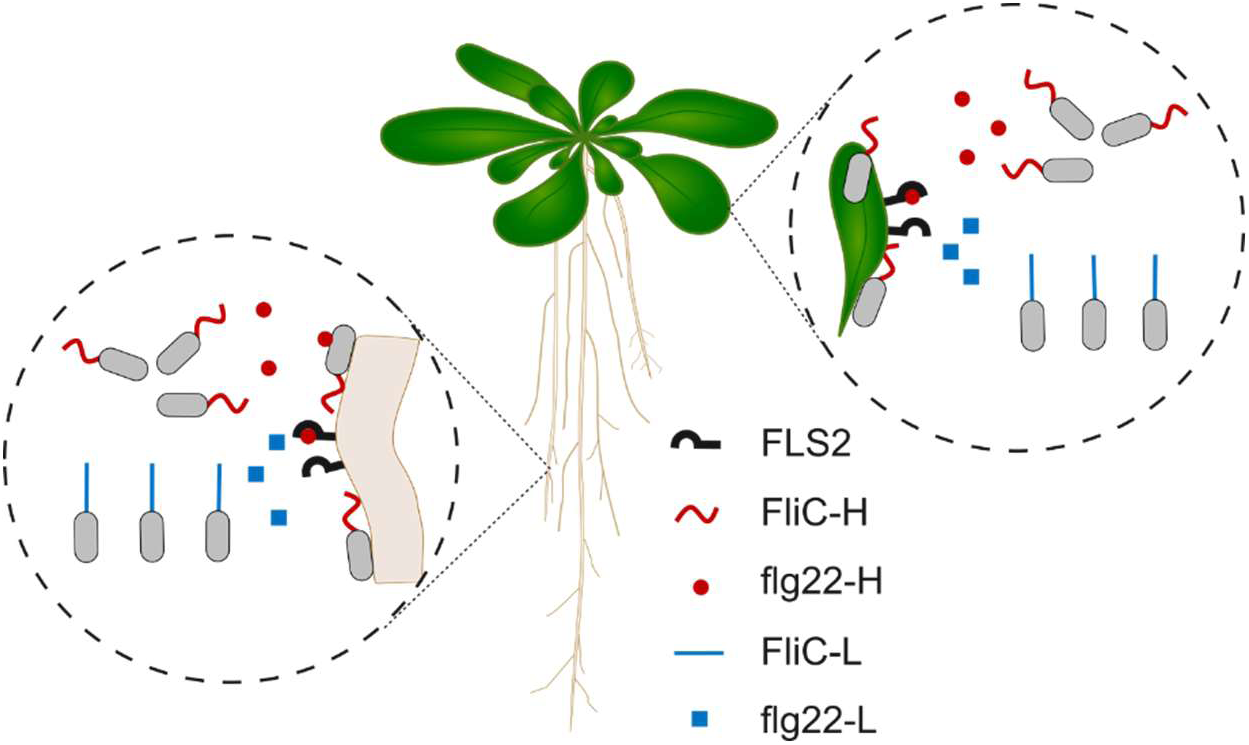
Model: The plant immune receptor FLS2 specifically targets *Sphingomonas* sp. *FliC* variants required for full plant colonization. Flagella of Sphingomonas MF220 contribute to swimming motility, plant colonization, and attachment to plant roots. While flagella composed of the non-immunogenic FliC-L protein (blue) facilitates more efficient swimming up a nutrient gradient, flagella containing the immunogenic FliC-H protein (red) are more effective at colonizing both the shoot and the root, and at attaching to plant roots. To effectively monitor bacteria capable of efficient root attachment and plant colonization, the plant FLS2 immune receptor has evolved to specifically recognize flg22-H epitopes (red triangles, derived from the immunogenic FliC-H required for attachment), rather than the flg22-L epitope (blue squares, derived from the non-immunogenic FliC-L essential for directional swimming).

Flagellin-FLS2 interactions are typically framed through the lens of antagonistic pleiotropy that balances immune recognition and motility (12). This model looks at mutational variations in a single flagellin gene performing multiple functions and predicts that immune evasion cannot occur without fitness costs. We examined attachment and colonization in *Sphingomonas*, where independently derived and regulated flagella provide disparate functions. Our findings reveal an alternative evolutionary trajectory for flagellar diversification. Instead of optimizing a single flagellin for immune evasion, motility, and colonization, *Sphingomonas* resolves conflict with the host plant via a division of labor. In certain members of this genus, a highly divergent collection of motility-flagella are immune-evasive while the flagella required for full colonization inside plant organs are immunogenic. In that case, even if the more ancient FliC-H was responsible for both motility and attachment, the acquisition of the motility-only FliC-L could enable a possible dual function flagella to specialize in attachment, and lose it’s motility function. Thus, *Sphingomonas* sp. avoids the pleiotropic constraint identified by Parys et al. (2021) (12) by separating motility and attachment to different flagellar system rather than by altering a constrained epitope.

Spatial receptor distribution provides a mechanistic context for the flagellar specialization we describe in *Sphingomonas* sp.. In Arabidopsis, FLS2 is not uniformly present across the plant body. Instead, FLS2 accumulates in potential microbial entry or attachment points inside the shoot and in compartments protected by the endodermal barrier in roots. (28, 29). This spatial organization suggests that immune surveillance is most intense where microbes transition from swimming and exploring to intimate host contact. Our data showing that FliC-H is required for colonization and expressed to a higher degree in colonizing bacteria, compared to host organ surface bacteria, suggest that the spatial organization of FLS2 is aimed at detecting FliC-H where it is most needed. Thus, receptor localization on the host and flagellin deployment by the bacterium converge to structure the outcome of immune perception.

The evolutionary strategy we define may lead to immune-mediated détente between the host and strains of *Sphingomonas* carrying divergent *FliC* loci: some *Sphingomonas* sp. are allowed to colonize plant organ surfaces via FliC-L driven flagellar motility but are gated out of shoot and root endophytic compartments by the requirement of FliC-H for attachment and colonization. This model supports the concept that the plant benefits provided by various *Sphingomonas* sp. might be largely limited to non-endodermal compartments by the FLS2-mediated host immune response in endodermal compartments. The weak MTI response generated by at least one *Sphingomonas* sp. might reflect this gating in action as the bacteria are continually probing the plant’s inner niches and thus triggering weak MTI (31, 32).

Sphingomonas are not the only bacteria encoding more than one flagellum, yet the specialization and division of labor we identified for *Sphingomonas* is unique. In biphasic *Salmonella enterica*, phase variation between two distinctly immunogenic flagellins (FliC and FljB) facilitates immune evasion in animal hosts (35), but the two proteins are functionally redundant for motility. Recent work in plant-associated *Pseudomonas* identified strains with duplicated *FliC* genes that differ in immunogenicity (36), though only one copy supports flagellar biogenesis and motility while the second is poorly expressed and non-functional. Importantly, *Sphingomonas* sp. strain A1 also possesses multiple distinct flagellin genes that assemble into a single heteromeric flagellum (37, 38). In this strain, the flagellum is a bifunctional appendage required for both motility and alginate uptake (39, 40). The alginate binding properties of *Sphingomonas* A1 flagella provide a plausible precedent for our observations in Sphingomonas MF220, where the FliC-H variant may have evolved a specialized role in promoting attachment to plant tissues, independent of the flagellum’s primary role in motility. Other genera, including *Vibrio* and *Aeromonas*, use physically distinct polar and lateral flagella for different types of motility (41). In contrast, our results show functional specialization across two flagellins, suggesting division of labor strategy, where one enables full directional swimming and immune evasion, and the other supports colonization despite being immunogenic.

In summary, we propose a refined model of MAMP-PRR coevolution: instead of targeting traits required for generic microbial viability, like swimming, plant immune receptors preferentially target bacterial functions directly associated with intimate host interaction, like attachment and colonization. This turns the plant immune system into a strategic gatekeeper in commensal interactions, enabling microbial arrival and spatially limited proliferation but restricting the transition from surface exploration to persistent colonization and damaging bacterial population expansion.

## Material and Methods

### Synteny analysis of the genomic island surrounding the FliC genes

To examine genomic context and conservation of flagellum-associated regions, we performed a synteny analysis centered around each *FliC* gene. For every genome, we extracted all genes located upstream and downstream of both *FliC* loci and annotated them using KEGG Orthology (KO) identifiers. Genes assigned to flagellum-related KOs (e.g., motor, basal body, hook, export apparatus) were used to define a “flagellum island,” operationally defined as the continuous genomic segment containing *FliC* and its adjacent flagellar genes. We then compared the flagellum islands surrounding the 12 focal *FliC* alleles at two levels: (i) gene content similarity, quantified using the Jaccard index based on shared KO identifiers, and (ii) gene order conservation, quantified as the fraction of conserved adjacent gene pairs (i.e., the proportion of neighboring genes in island A that remain neighbors in island B). For each pairwise comparison, we calculated an island similarity index by multiplying the gene-content Jaccard index and the adjacency index, providing a composite measure of synteny that captures both presence/absence of genes and conservation of local genomic architecture.

### Identification of *FliC* genes and cognate flg22 epitopes in wild *Sphingomonas*

To identify FliC homologs in environmental *Sphingomonas* isolates, we constructed a hidden Markov model (HMM) profile from the 12 experimentally flg22-validated FliC proteins in our strain collection and searched each genome using HMMER (42). Hits with high statistical confidence (E-value < 1×10⁻⁵⁰) were retained as bona fide FliC candidates.

Next, to extract the corresponding flg22 epitope from each identified FliC gene, we developed a guided positional alignment strategy. We started by identifying alignment consensus sequences built exclusively from the six strains for which flg22 immunogenicity had been experimentally characterized. We aligned each feature (FliC-H, FliC-L, flg22-H, flg22-L) independently. To construct four alignment consensus sequences. Then, for every newly identified FliC sequence, we performed a pairwise alignment to the FliC-H, FliC-L, flg22-H, and flg22-L consensus sequences to identify the correct region of the protein. The 22-amino-acid segment mapping to the flg22 position was extracted as the predicted flg22 sequence. This two-step positional alignment allowed us to reliably identify the flg22 epitope from highly divergent FliC and flg22 sequences, ensuring accurate recovery of the correct region across hundreds of wild *Sphingomonas* genomes.

### Constructing a *Sphingomonas* MLST tree

To infer phylogenetic relationships among *Sphingomonas* isolates, we constructed a multilocus sequence typing (MLST) tree based on conserved core genes. We identified 48 single-copy housekeeping genes present across all genomes in the dataset. Each gene was aligned independently using MAFFT (43). The resulting alignments were concatenated into a single “core-genes-genome”. To reduce noise from poorly aligned or rapidly evolving regions, we applied ZORRO (44) to compute highly divergent regions in the alignment. Finally, a maximum-likelihood phylogenetic tree was then generated using FastTree2 (45).

### Swimming assay

*Sphingomonas* strains MF220_WT_, MF220_DD,_ MF220_L_ and MF220_H_ were grown overnight at 28 °C in 2×YT medium supplemented with rifampicin (50 µg/mL). The following day, 1 mL of each culture was centrifuged at 6,000 × g for 5 minutes, washed twice with fresh 2×YT medium, and resuspended in 500 µL of 2×YT. For motility assays, 2 µL of each bacterial suspension was spotted onto 0.3% LB soft agar plates and allowed to dry. Plates were incubated at 28 °C for 48 hours, after which swimming halos were imaged, and the radius of the swimming zone was used to quantify motility. At the end of swimming experiments, bacteria were sampled from the outermost swimming halo and from the center of the swimming MF220_WT_ population for expression analysis (Methods). In addition, bacteria were also collected from a 2-days old bacterial colony grown on a standard 1.5% LB agar plate as reference.

### Deleting *FliC*-L and *FliC*-H from Sphingomonas MF220

We generated three *Sphingomonas* MF220 deletion derivatives lacking either *FliC-L* (MF220_H_ derivative), *FliC*H (MF220_L_ derivative), or both genes (MF220_DD_ derivative) using homologous recombination. To construct the deletion plasmids, we amplified ∼1,000 bp regions upstream and downstream of each *FliC* gene from MF220 genomic DNA. For MF220_L_ derivative, we used primers GD03_flkU_FliCH_F/GD04_flkU_FliCH_R and GD05_flkD_FliCH_F/ GD06_flkD_FliCH_R; for MF220_H_ derivative, we used GD11_flkU_FliCL_F/ GD12_flkU_FliCL_R and GD13_flkD_FliCL_F/GD14_flkD_FliCL_R. The internal primers (GD04_flkU_FliCH_R and GD05_flkD_FliCH_F, and their MF220_H_ derivative counterparts) included 5’ overhangs complementary to one another, while the outer primers contained 5’ overhangs homologous to the linearized pMo130 vector backbone.

We assembled the flanking fragments and pMo130 by Gibson assembly to generate pMo130_dFliCH and pMo130_dFliCL. We transformed the constructs into *E. coli* Top10 via heat shock and selected transformants on LB agar containing 50 µg/mL kanamycin. We validated colonies by PCR using flanking primers GD07_Vo_FliCH_F/GD08_Vo_FliCH_R and GD15_Vo_FliCL_F/ GD16_Vo_FliCL_R, propagated the plasmids in 2×YT medium with 50 µg/mL kanamycin, and purified them using the QIAprep Spin Miniprep Kit (QIAGEN).

We electroporated pMo130_dFliCH and pMo130_dFliCL into *E. coli* WM3064, a diaminopimelate (DAP)-auxotrophic conjugation donor strain. We selected transformants on LB agar supplemented with 0.3 mM DAP, 50 µg/mL kanamycin, and 50 µg/mL erythromycin. We mated the resulting donor strains with MF220 wild-type cells on LB agar with 0.3 mM DAP. After overnight incubation, we selected transconjugants on LB agar with 50 µg/mL kanamycin.

We purified single crossover transconjugants by re-streaking and counter selected for double recombinants on LB agar containing 10% sucrose. We screened sucrose-resistant colonies by PCR: primers GD07_Vo_FliCH_F/GD08_Vo_FliCH_R (for MF220_L_ derivative) or GD15_Vo_FliCL_F/ GD16_Vo_FliCL_R (for MF220_H_ derivative) amplified across the flanking regions and yielded a product only upon deletion, while primers GD09_Vi_FliCH_F/GD10_Vi_FliCH_R (for MF220_L_ derivative) or GD17_Vi_FliCL_F/ GD18_Vi_FliCL_R (for MF220_H_ derivative) within the deleted gene yielded no product. We confirmed deletion in MF220_H_ derivative and MF220_L_ derivative and further generated the MF220_DD_ strain by sequential deletion in the MF220_H_ derivative background using the same strategy.

All oligonucleotide sequences used are listed in the SI Appendix, Table S2.

### Scanning Electron Microscopy (SEM)

*Sphingomonas* strains used in this study included MF220_WT_, MF220_DD,_ MF220_L_ and MF220_H_. Cultures were grown overnight at 28 °C in 2×YT medium supplemented with rifampicin (50 µg/mL), then diluted 1:100 into fresh 2×YT medium (without antibiotics) and incubated at 28 °C with shaking until reaching mid-log phase (OD₆₀₀ ≈ 0.4–0.5). To preserve intact flagella, cells were harvested by gentle centrifugation at 1,000 × g for 2 minutes at room temperature, then immediately resuspended in fixative solution (2% paraformaldehyde/2.5% glutaraldehyde in 0.15 M sodium phosphate buffer, pH 7.4) for subsequent processing and imaging by scanning electron microscopy. A droplet of each fixed sample was plated on 12 mm round glass coverslips and allowed to adhere at 4°C overnight. Coverslips were washed three times with 0.15 M sodium phosphate buffer, then incubated with 1% buffered osmium tetroxide for 30 minutes at room temperature. Coverslips were then washed three times in deionized water and dehydrated through an ascending series of ethanol (30%, 50%, 75%, 100%, 100% 100%). Coverslips were transferred in 100% ethanol to a Samdri-795 critical point dryer and dried using liquid carbon dioxide as the transitional solvent (Tousimis Research Corporation, Rockville, MD). Once dry, coverslips were mounted on 13 mm diameter aluminum stubs using carbon adhesive tabs and sputter coated with 5 nm of a 60:40 gold/palladium alloy using a Cressington 208HR Sputter Coater (Ted Pella Inc., Redding, CA). Images were obtained using a Zeiss Supra 25 FESEM operating at 5 kV using an SE2 detector, 20 µm aperture, and approximate working distances of 5 to 8 mm (Carl Zeiss Microscopy, LLC, Peabody, MA).

### Characterizing short-term attachment to the plant

Three-week-old *A. thaliana* Col-0 plants grown on 0.5× MS agar plates were used for bacterial attachment assays. Shoots and roots were carefully separated and transferred into individual wells of a sterile 12-well plate. Plant tissues were submerged in 2 mL of bacterial suspension (10⁶ CFU/mL in sterile distilled water) containing *Sphingomonas* strains: MF220_WT_, MF220_DD,_ MF220_L_ and MF220_H_. Inoculated plates were incubated with subtle shaking at 50 rpm for 4 hours. After incubation, tissues were washed three times with sterile water and vortexed briefly between washes to remove loosely attached bacteria. Washed tissues were weighed, transferred to microcentrifuge tubes containing three 4-mm beads and 250 μL of sterile water, and then homogenized using a bead mill. Serial dilutions were spotted onto LB agar supplemented with rifampicin (50 μg/mL), and colony-forming units (CFUs) were counted after 48 hours of incubation at 28°C. Attachment was quantified as CFUs per mg of fresh tissue. Experiments were performed with at least three independent biological replicates per strain and tissue type.

### Characterizing long-term plant colonization. Root Colonization

Surface-sterilized *A. thaliana* seeds (Col-0 background, *fls2* mutant) were stratified at 4°C in the dark for 2 days and then transferred to 0.5× MS agar plates to germinate under long-day conditions (16 h light / 8 h dark) at 22°C. Seven-day-old seedlings were carefully transferred to new 0.5× MS plates supplemented with *Sphingomonas* strains (MF220_WT_, MF220_DD,_ MF220_L_ and MF220_H_) adjusted to ∼10⁷ CFU/plate. After 12 days of co-cultivation, roots were harvested and washed three times with sterile distilled water to remove loosely attached bacteria. Roots were then weighed, ground in water, and serial dilutions were plated on 2X YT agar plates containing Rifampicin (50 µg/ml). CFUs were enumerated after 48 hours of incubation at 28°C. Colonization data were normalized to root weight. Each experiment was performed with at least three biological replicates.

### Shoot Colonization

Four-to six-week-old *A. thaliana* plants (Col-0 and *fls2*) grown in pots were spray-inoculated with bacterial suspensions (10⁹ CFU/mL) of *Sphingomonas* MF220 strains ( MF220_WT_, MF220_DD,_ MF220_L_ and MF220_H_). Plants were maintained under short-day conditions for up to 12 days post-inoculation (dpi). Leaves were harvested at 0, 3, 6, 9, and 12 dpi, and the fresh weight of each sample was recorded prior to bacterial recovery. For each strain, three leaves were pooled per tube, with three independent tubes used per experimental set. To recover bacteria, leaves were washed three times in 1 mL sterile water for 30 s each. The first wash fraction was retained and designated as the loosely attached epiphytic population. Washed leaves were then transferred to 2 mL tubes containing three 4-mm glass beads and 500 µL sterile water and homogenized using a bead mill to release tightly attached and endophytic bacteria. Serial dilutions of all fractions were prepared in 96-well plates and spotted on selective medium for CFU enumeration after 2 days of incubation at 28°C.

### Tissue Harvest for In Planta Characterization of *FliC* expression

For expression analysis, shoot and root tissues were harvested separately from a colonization assay. Surface-sterilized *A. thaliana* Col-0 seeds were stratified at 4°C in the dark for 2 days, then transferred to 0.5× MS agar plates to germinate under long-day conditions (16 h light / 8 h dark) at 22°C. Seven-day-old seedlings were carefully transferred to new 0.5× MS plates supplemented with 10⁷ CFU/plate MF220_WT_ *Sphingomonas*. After 12 days of co-cultivation under long-day conditions, roots were harvested and washed three times with sterile distilled water to remove loosely attached bacteria. The first wash was processed as loosely attached bacteria (root rhizosphere). After three washes, roots were frozen in liquid nitrogen to process firmly attached rhizoplane and endophytic bacteria. Similarly, for shoots, the first water wash was processed as loosely attached bacteria (leaf phyllosphere). Subsequently, surface-sterilized shoots by immersion in 70% ethanol for 60 seconds, followed by two rinses with sterile distilled water. Finally, shoots were frozen in liquid nitrogen for endophytic bacterial gene expression analysis. Each experiment was performed in three biological replicates.

### RNA Isolation and qRT-PCR

Total RNA was extracted using the ZymoBIOMICS DNA/RNA Miniprep Kit (Zymo Research, Cat. R2002) following the manufacturer’s instructions. The isolated RNA was treated with DNase I to remove any DNA contamination using the Turbo DNA Free Kit (Invitrogen, Cat. AM1907). cDNA synthesis was performed using SuperScript IV Reverse Transcriptase (Invitrogen). Quantitative real-time PCR (qRT-PCR) was carried out using PowerUp SYBR Green Master Mix (Applied Biosystems) on a ViiA™ 7 Real-Time PCR System (Applied Biosystems). qRT-PCR was performed with an initial denaturation at 95 °C for 10 min, followed by 40 cycles of 95 °C for 20 s, 58 °C for 20 s, and 72 °C for 30 s, with fluorescence acquisition during the extension step. Melt-curve analysis was performed using the instrument’s default settings to confirm amplification specificity. Expression levels of *FliC-*H and *FliC-*L were calculated based on –ΔCT values normalized to *recA* as the internal control. No amplification was detected in the no-bacteria negative control.

All oligonucleotide sequences used are listed in the SI Appendix, Table S2.

## Supporting information

Supplementary Apendix

## Acknowledgements

This work was supported by NSF grant IOS-2416244 to JLD and by the Howard Hughes Medical Institute. DR gratefully acknowledges funding from a EMBO Long Term Fellowship (ALTF 743–2019). CRF gratefully acknowledges funding from a Natural Sciences and Engineering Research Council of Canada postdoctoral fellowship (532852-2019). DSL is supported by a Wallenberg Academy Fellows grant (https://kaw.wallenberg.org/en, 2021.0102). This work was supported by HHMI. Scanning Electron Microscopy was performed at the Microscopy Services Laboratory, Department of Pathology and Laboratory Medicine, University of North Carolina at Chapel Hill, which is supported in part by P30 CA016086 Cancer Center Core Support Grant to the UNC Lineberger Comprehensive Cancer Center. We thank Kristen White, Jillann Madren, and Victoria Madden for expert technical assistance and microscopy support. We thank Prof. Sarah R. Grant for critical reading and all members of the Grant-Dangl lab for helpful discussions during this work.

