## Supplementary Apendix for "Specialization of independently acquired flagellar FliC proteins in plant-associated *Sphingomonas* balances swimming and immunogenicity"

**This PDF file includes:**

Figures S1 to S8
Table S1 to S2
Legends for Movie S1

**Other supporting materials for this manuscript include the following:**

Movie S1

**Figures**

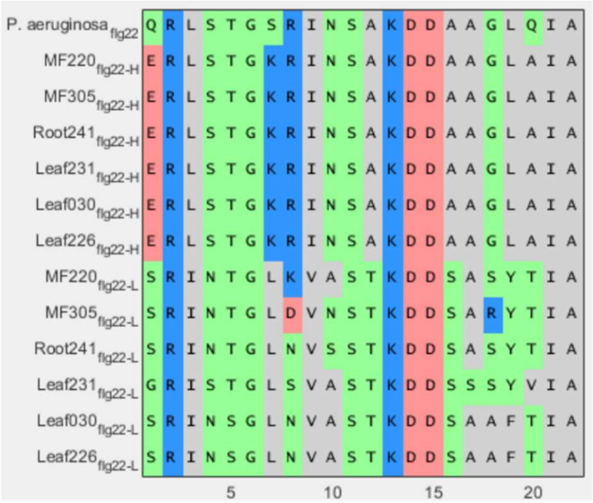


**Fig. S1. Sequences of functionally tested flg22 epitopes from six *Sphingomonas* isolates with diverging *FliC* genes.**

The peptide sequence of the six immunogenic (flg22-H) and six non-immunogenic (flg22-L) epitopes from *A. thaliana*-associated Sphingomonas carrying both immunogenic and non-immunogenic *FliC* genes. The canonical flg22 sequence from *P. aeruginosa* is included at the top for reference.

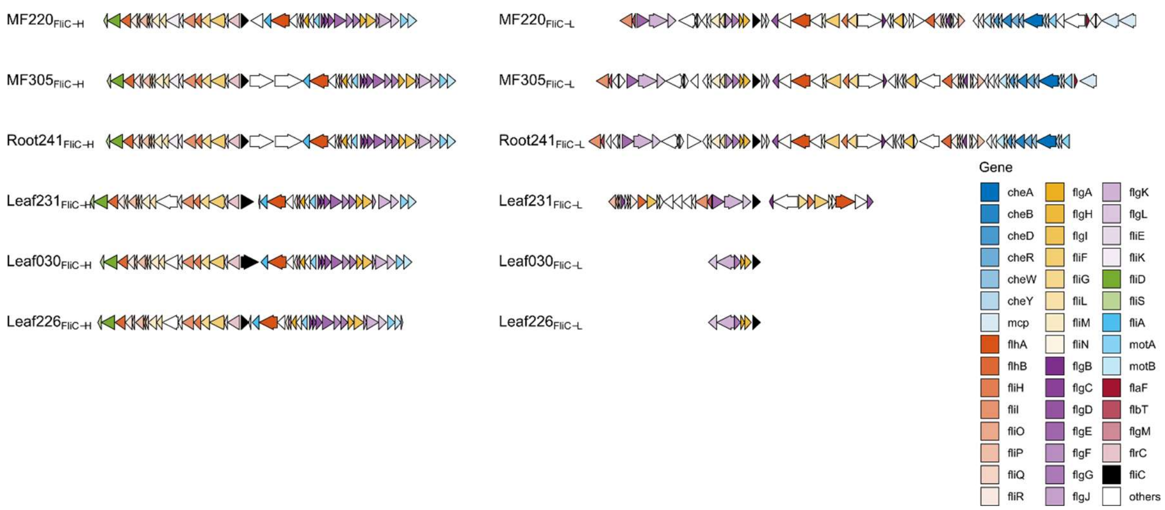


**Fig. S2. The genomic island encompassing the immunogenic *FliC-H* lacks chemotactic machinery.**

The organization of genomic islands containing *FliC*-H (left panel) and *FliC*-L (right panel) in *Sphingomonas* isolates is depicted. Within each island, the respective *FliC* gene is highlighted in black, and other flagella-related genes are color-coded by functional category. Notably, while three of the six *FliC*-L islands contain genes encoding chemotaxis machinery (shades of blue) while none of the *FliC*-H islands possess these genes.

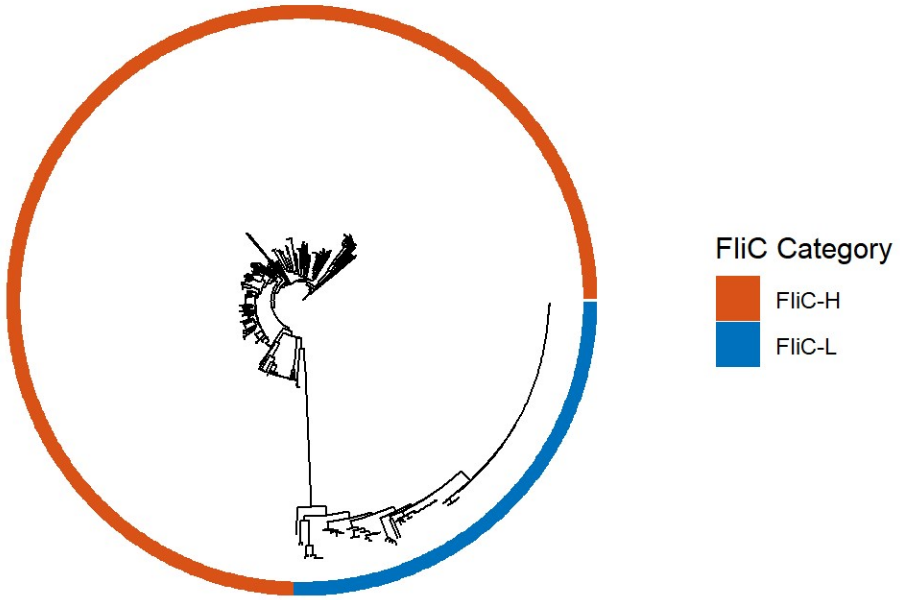


**Fig. S3. Phylogenetic distribution of FliC proteins corresponds to flg22-predicted immunogenicity.**

Maximum-likelihood gene tree of all FliC protein sequences identified across a 400-member Sphingomonas isolate collection. The outer ring indicates predicted immunogenicity based on the specific flg22 epitope sequence carried by each gene. The phylogeny demonstrates that the evolutionary history of the full-length FliC mirrors the functional divergence of its flg22 epitope, with distinct clades corresponding to predicted immunogenic and non-immunogenic variants.

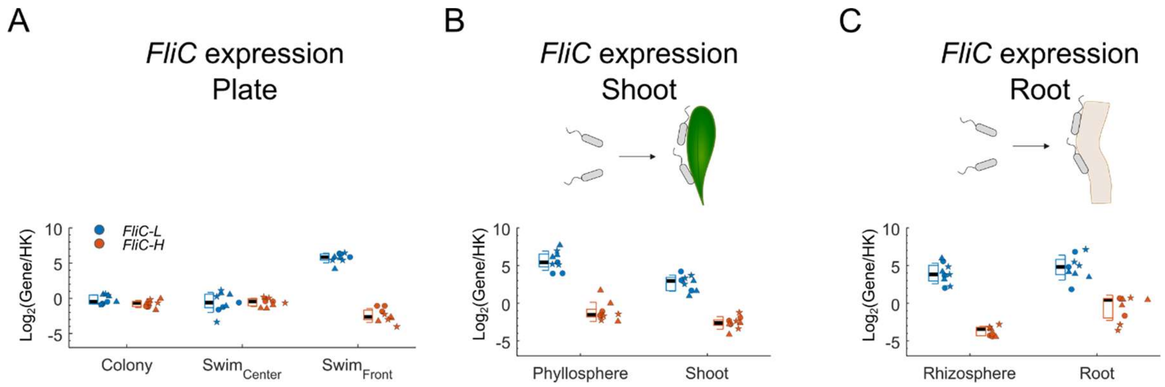


**Fig. S4. Expression of the two *FliC* genes**

Expression of *FliC*-H (red) and *FliC*-L (blue) in MF220 from a bacterial in swimming assay (A) and in-planta after 12 days colonizing A. thaliana shoot (B) and root (C). **(A)** Fast-swimming bacteria at the front of a swimming plate express more *FliC*-L and slightly less *FliC*-H than non-swimming bacteria in colony and from slow-swimming bacteria in the center of a swimming population (n = 9 biologically independent samples from 3 independent experiments, denoted by shape). **(B)** Shoot-associated bacteria (right) exhibit lower *FliC*-H and *FliC*-L expression than non-attached bacteria in the phyllosphere (linear model, $P < 0.05$). Since *FliC*-H only showed a modest effect and *FliC*-L showed a stronger effect, overall, the ratio of *FliC*-H to *FliC*-L expression shifts towards *FliC*-H upon shoot colonization (Fig. 4C top; n = 9 biologically independent samples from 3 independent experiments, denoted by shape). **(C)** Root-associated bacteria (right, comprising rhizoplane and endophytes) exhibit significantly higher *FliC*-H expression than non-attached bacteria in the rhizosphere (left; linear model, $P < 0.05$). In contrast, *FliC*-L expression remains unchanged. Overall, the *FliC*-L to *FliC*-H ratio shifts to *FliC*-H upon colonization (Fig. 4C bottom; n = 9 biologically independent samples from 3 independent experiments, denoted by shape). In all panels, boxplots show the median (black line), 1st and 3rd quartiles (box), and one standard deviation (whiskers) of treatment.

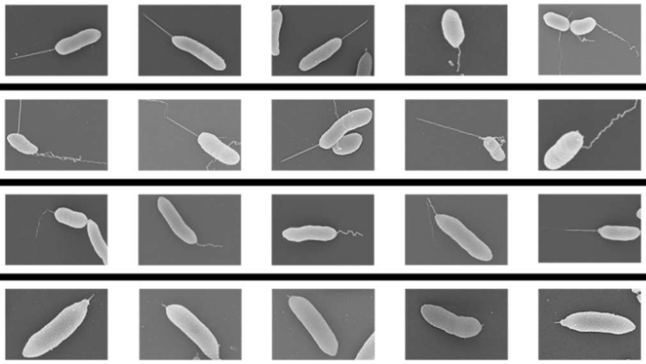

| | Flagella: Straight | Flagella: Wobbly |
|---|---|---|
| MF220$_{WT}$ | 28 | 3 |
| MF220$_{L}$ | 16 | 3 |
| MF220$_{H}$ | 11 | 26 |
| MF220$_{DD}$ | 0 | 0 |

**Fig. S5. The two flagellar variants for distinct morphologies.**

Five representative scanning electron microscopy images of the four MF220 derivatives (left) and quantification of the two distinct morphologies in each derivative (right). Both the MF220$_{WT}$ and the MF220$_{L}$ variant are dominated by straight spear like flagella, while the MF220$_{H}$ is dominated by a wobbly, twisted flagella. None of the double deletion MF220$_{DD}$ bacteria caried any flagella.

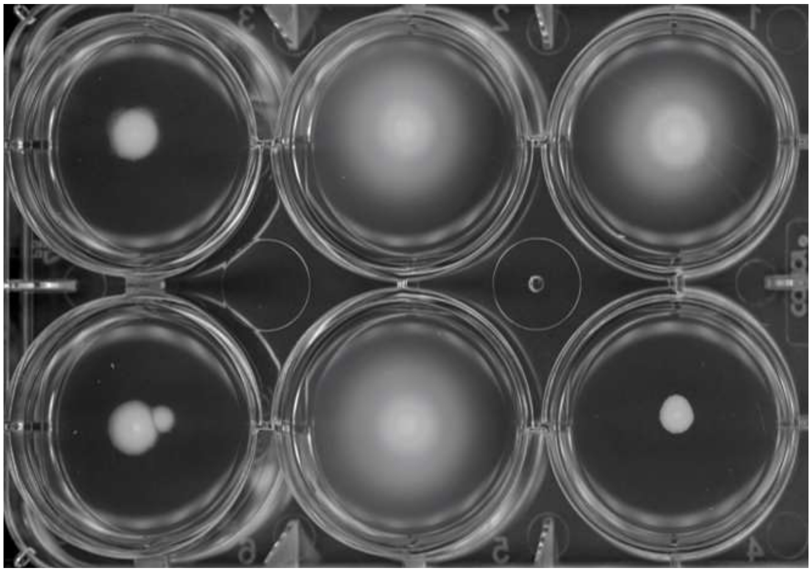


**Fig. S6. FliC-L is essential for full directional swimming motility.**

Representative images of swimming populations for *Sphingomonas* MF220 derivatives after three days of incubation in semi-solid agar. The $MF220_L$ derivative (blue, two independent deletion alleles), carrying only the non-immunogenic *FliC*-L gene, exhibits swimming motility comparable to that of the wild-type $MF220_{WT}$ (gray). In contrast, the $MF220_H$ derivative (red, two independent deletion alleles), lacking *FliC*-L, is severely compromised in swimming. The double deletion derivative ($MF220_{DD}$, purple), which lacks both *FliC* genes, shows even further reduced swimming compared to the $MF220_H$ derivative.

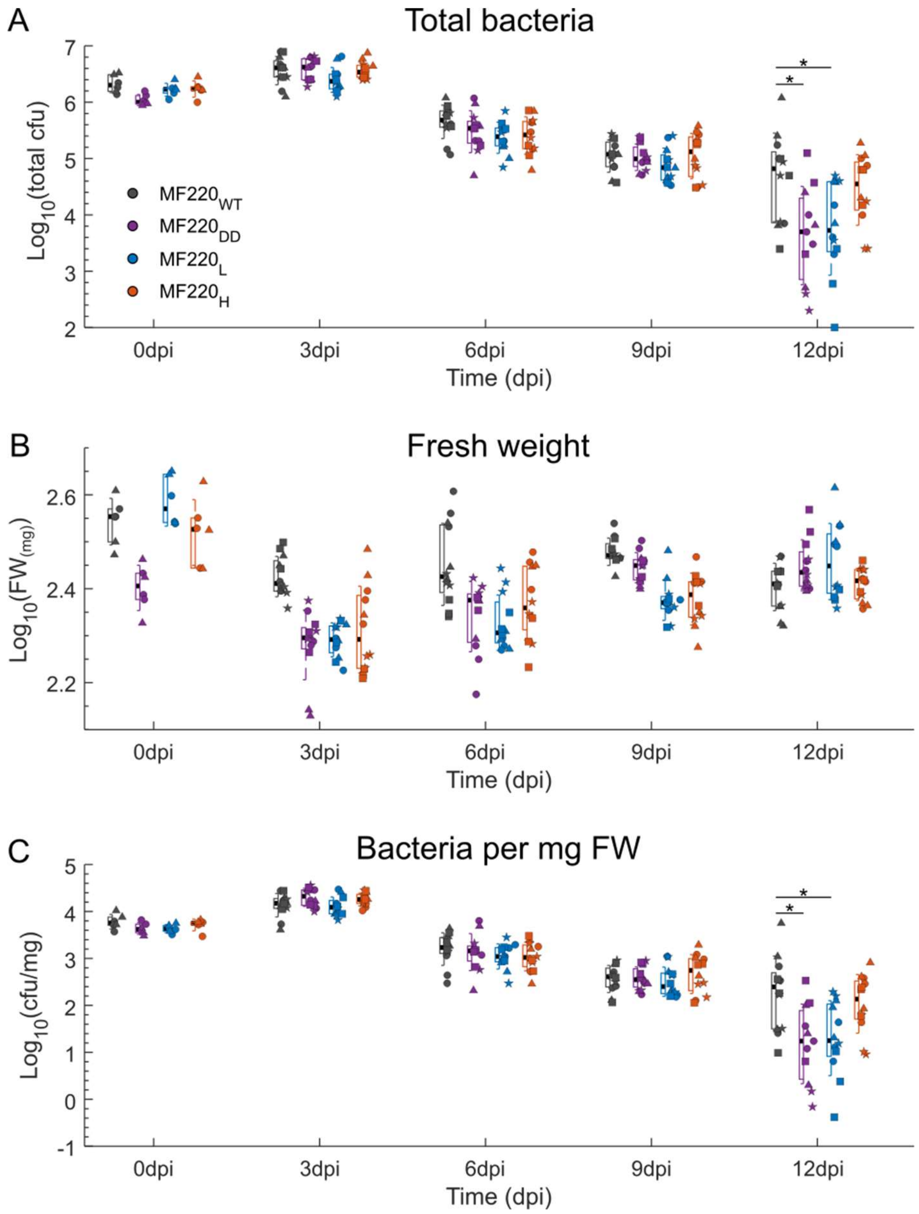


**Fig. S7. The abundance of spray-inoculated MF220 on *A. thaliana* leaves declines over time.**

Qualification of the four MF220 derivatives spray inoculated on wild-type *A. thaliana* Col-0 leaves over time. The total bacterial load **(A)**, tissue fresh weight **(B)**, and bacterial load per mg fresh weight **(C)** are plotted for each timepoint. The bacterial load declines over time for all derivatives, but $MF220_{WT}$ and $MF220_{H}$ derivatives survive more than the $MF220_{DD}$ and $MF220_{L}$ (n = 9 biologically independent samples from 3 independent experiments, denoted by shape, except for Day 0, where n = 6 biologically independent samples from two independent experiments). The biggest effect is seen at 12 dpi (Fig. 4B top). Boxplots show the median (black line), 1st and 3rd quartiles (box), and one standard deviation (whiskers) of treatment. Asterisk indicates biological and statistical significance based on a linear model (P-value < 0.05 and $Log_{10}$(Fold change) > 0.5; comparisons without markings are nonsignificant).

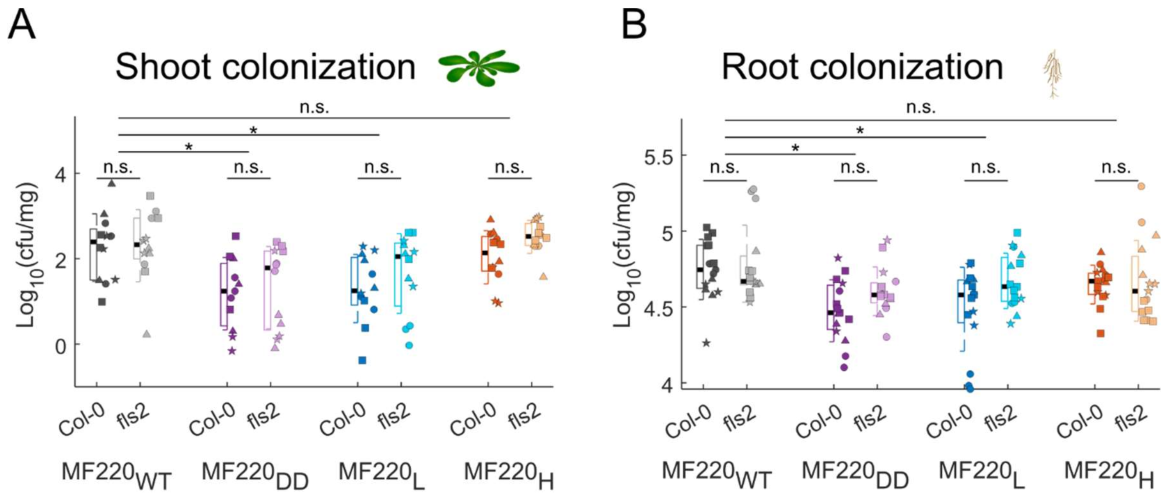


**Fig. S8. The contribution of FliC-H to MF220 colonization is FLS2-independent.**

Qualification of the four MF220 derivatives colonizing the shoots (A) and roots (B) of wild type *A. thaliana* and the *fls2* mutant after 12 days. To test if the colonization phenotypes observed were dependent on the host immune receptor FLS2 we analyzed the log transformed bacterial load ($Log_{10}$(cfu/mg)) using a linear mixed effect model with an interaction term: $Log_{10}$(cfu/mg) ~ Host + Bacteria + Host*Bacteria + (1|Exp) where, Host is either Col-0 or *fls2*, Bacteria represents the four MF220 derivatives, Host*Bacteria is the interaction effect and (1|Exp) is the random intercept accounting for variability between independent experiments. In both the shoots and the roots $MF220_{DD}$ (purple) and $MF220_{L}$ (blue) colonized the plant to a lesser degree than the wild-type $MF220_{WT}$ (gray). Conversely, the $MF220_{L}$ derivative (red) colonized at levels comparable to the wild type (linear model, $P < 0.05$, $n = 15$ biologically independent samples from 4 independent experiments, denoted by shape). An FLS2 dependence of the FliC derived plant-bacteria interactions would have manifested as a statistically significant Host*Bacteria interaction term. The lack of statistical significance of the interaction term (n.s.) indicating that the specific contribution of FliC-H to plant colonization is independent of FLS2. In all panels, boxplots show the median (black line), 1st and 3rd quartiles (box), and one standard deviation (whiskers) of treatment.

**Tables**

**Table S1. FliC genes of the six Sphingomonas identified as carrying diverging flg22 epitpes**

| Isolate | Strain_ID | FliC-H (gene_ID) | FliC-L (gene_ID) |
|---|---|---|---|
| Sphingomonas sp. MF220 | 2643221532 | 2643690330 | 2643691427 |
| Sphingomonas sp. MF305 | 2565956511 | 2565998623 | 2565996948 |
| Sphingomonas sp. Root241 | 2643221622 | 2644127311 | 2644128520 |
| Sphingomonas sp. Leaf231 | 2643221782 | 2644931960 | 2644933920 |
| Sphingomonas sp. Leaf030 | 2643221763 | 2644849962 | 2644852765 |
| Sphingomonas sp. Leaf226 | 2643221757 | 2644829900 | 2644829262 |

**Table S2. Oligos used in the paper**

| Num | Name | Sequence | notes |
|---|---|---|---|
| 1 | RT01_FliC-H_F | GAGGACACCGACTTCTCGAC | Primers for qRT-PCR |
| 2 | RT02_FliC-H_R | AGAAGCGAAAGAACCGTCTG | |
| 3 | RT03_FliC-L_F | CTCCAGAGCCTGAACGAGAC | |
| 4 | RT04_FliC-L_R | TGAGCAATGGTGTACGAAGC | |
| 5 | RT05_recA_F | CAACCAGCTGCGCATGAAGA | |
| 6 | RT06_recA_R | GAGGCGTAGAACTTGAGCGC | |
| 7 | GD01_pMo130 F | tgcagcggatccctctag | Primers for a clean deletion of the FliC-H, FliC-L and Both from *Sphingomonas* MF220 |
| 8 | GD02_pMo130 R | Gctcactcaaaggcggtaatac | |
| 9 | GD03_flkU_FliC-H_F | catgcatctagagggatccgctgcaTGACTTCGCCCTGCAGATC | |
| 10 | GD04_flkU_FliC-H_R | cttatcggagaagcgaaagaGTTCGTTACTCCATGCTCGTC | |
| 11 | GD05_flkD_FliC-H_F | gacgagcatggagtaacgaacTCTTTCGCTTCTCCGATAAGC | |
| 12 | GD06_flkD_FliC-H_R | accgtattaccgcctttgagtgagcTATCACCACCAATGCGATCG | |
| 13 | GD07_Vo_FliC-H_F | AATTGCCCATTTCGAGACCG | |
| 14 | GD08_Vo_FliC-H_R | CCTCTCGAACGACATTGCAA | |
| 15 | GD09_Vi_FliC-H_F | GAGGACACCGACTTCTCGAC | |
| 16 | GD10_Vi_FliC-H_R | AGAAGCGAAAGAACCGTCTG | |
| 17 | GD11_flkU_FliC-L_F | accgtattaccgcctttgagtgagcGTCCACTTGCAGCTTACAGG | |
| 18 | GD12_flkU_FliC-L_R | ggccatagactccagggaaaGTACGCCCGCATGTTCATT | |
| 19 | GD13_flkD_FliC-L_F | aatgaacatgcgggcgtacTTTCCCTGGAGTCTATGGCC | |
| 20 | GD14_flkD_FliC-L_R | catgcatctagagggatccgctgcaCAAGTTCGGGAAGATCAGCC | |
| 21 | GD15_Vo_FliC-L_F | GCCACATCGACATGAGGATC | |
| 22 | GD16_Vo_FliC-L_R | CGTCAACCACTATAGACGCG | |
| 23 | GD17_Vi_FliC-L_F | CTCCAGAGCCTGAACGAGAC | |
| 24 | GD18_Vi_FliC-L_R | TGAGCAATGGTGTACGAAGC | |

**Movie S1 (separate file). Swimming dynamics of MF220 derivatives on semi-solid agar.**

The video captures the swimming migration of four MF220 derivatives: $MF220_{WT}$ (gray), $MF220_{DD}$ (purple), $MF220_{L}$ (blue), and $MF220_{H}$ (red). The area of the bacterial population was monitored and recorded hourly over a three-day period to quantify their migration dynamics.
